# Long-term nutrient enrichment of an oligotroph-dominated wetland increases bacterial diversity in bulk soils and plant rhizospheres

**DOI:** 10.1101/2020.01.08.899781

**Authors:** Regina B. Bledsoe, Carol Goodwillie, Ariane L. Peralta

## Abstract

In nutrient-limited conditions, plants rely on rhizosphere microbial members to facilitate nutrient acquisition, and in return plants provide carbon resources to these root-associated microorganisms. However, atmospheric nutrient deposition can affect plant-microbe relationships by changing soil bacterial composition and by reducing cooperation between microbial taxa and plants. To examine how long-term nutrient addition shapes rhizosphere community composition, we compared traits associated with bacterial (fast growing copiotrophs, slow growing oligotrophs) and plant (C3 forb, C4 grass) communities residing in a nutrient poor wetland ecosystem. Results revealed that oligotrophic taxa dominated soil bacterial communities and that fertilization increased the presence of oligotrophs in bulk and rhizosphere communities. Additionally, bacterial species diversity was greatest in fertilized soils, particularly in bulk soils. Nutrient enrichment (fertilized vs. unfertilized) and plant association (bulk vs. rhizosphere) determined bacterial community composition; bacterial community structure associated with plant functional group (grass vs. forb) was similar within treatments but differed between fertilization treatments. The core forb microbiome consisted of 602 unique taxa, and the core grass microbiome consisted of 372 unique taxa. Forb rhizospheres were enriched in potentially disease suppressive bacterial taxa and grass rhizospheres were enriched in bacterial taxa associated with complex carbon decomposition. Results from this study demonstrate that fertilization serves as a strong environmental filter on the soil microbiome, which leads to distinct rhizosphere communities and can shift plant effects on the rhizosphere microbiome. These taxonomic shifts within plant rhizospheres could have implications for plant health and ecosystem functions associated with carbon and nitrogen cycling.

**Importance:** Over the last century, humans have substantially altered nitrogen and phosphorus cycling. Use of synthetic fertilizer and burning of fossil fuels and biomass have increased nitrogen and phosphorous deposition, which results in unintended fertilization of historically low-nutrient ecosystems. With increased nutrient availability, plant biodiversity is expected to decline and bacterial communities are anticipated to increase in abundance of copiotrophic taxa. Here, we address how bacterial communities associated with different plant functional types (forb, grass) shift due to long-term nutrient enrichment. Unlike other studies, results revealed an increase in bacterial diversity, particularly, of oligotrophic bacteria in fertilized plots. We observed that nutrient addition strongly determines forb and grass rhizosphere composition, which could indicate different metabolic preferences in the bacterial communities. This study highlights how long-term fertilization of oligotroph-dominated wetlands could alter the metabolism of rhizosphere bacterial communities in unexpected ways.

## Introduction

The soil microbiome is critical for plant health, fitness, and diversity, especially in nutrient-limited environments (1–4). In particular, within the rhizosphere plants provide carbon (C) resources to soil microorganisms in exchange for nutrients such as nitrogen (N) and phosphorus (P). However, nutrient enrichment has been documented to disrupt plant-microbe mutualisms (2). Over the last century, agricultural fertilization and the burning of fossil fuels and biomass have indirectly led to nutrient deposition onto historically low-nutrient ecosystems (5–8). Nutrient enrichment generally causes reduced plant species diversity (9, 10) sometimes as a shift in plant functional types with an increase in grass biomass and loss of forb diversity (11–13). Fertilization has also been shown to decrease soil microbial diversity across cropland, grassland, forest, and tundra ecosystems (14–16). Despite patterns that have emerged from these bulk soil studies, it is less clear how changes in soil microbial diversity due to nutrient additions influence rhizosphere microbial community assembly and diversity. We address this knowledge gap by comparing changes in rhizosphere bacterial community composition of a grass and forb within a long-term fertilization experiment.

Both bulk soil matrix (i.e., not in contact with plant roots) properties and plant identity influence rhizosphere microbial communities. The bulk soil matrix is the reservoir of microbial diversity from which rhizosphere-associated microbial communities are selected; therefore, shifts in bulk soil microbial communities affect rhizosphere assemblages (17–19). In many cases N, N and P, and N-P-K fertilization decreases soil bacterial diversity (14–16). Additionally, nutrient enrichment selects for more copiotrophic (i.e., fast-growing, r-strategists) microbial heterotrophs that preferentially metabolize labile C sources versus oligotrophic (i.e., slow-growing, K-strategist) microbial species, which can metabolize complex C sources (20–23). A molecular marker to identify life history strategy (i.e., copiotroph or oligotroph) is rRNA *(rrn)* gene copy number (23–26). Bacterial taxa are estimated to contain 1-15 rRNA gene copies, with faster growing taxa containing higher gene copies than slower growing taxa (20, 23–27). Specifically, bacterial growth rate is limited by transcription rates of rRNA, such that growth rate is estimated to double with doubling of rRNA gene copy number Further, several studies indicate fertilization increases the abundance of copiotrophic bacterial groups within Actinobacteria, Alphaproteobacteria, and Gammaproteobacteria and decreases abundance in oligotrophic bacterial groups within Acidobacteria, Nitrospirae, Planctomycetes, and Deltaproteobacteria of bulk soils (15, 21, 28, 29). Additionally, copiotrophic taxa within Alpha-, Beta-, and Gamma-Proteobacteria, Actinobacteria, Firmicutes and Bacteroidetes are dominant members of some rhizosphere communities (17, 30, 31).

While the bulk soil environment is the primary source of rhizosphere diversity, plant species also influence rhizosphere bacterial community assembly due to variation in rhizodeposition (30–33). Rhizodeposits include nutrients, exudates, root cells, and mucilage released by plant roots (34). Plants allocate 5-20% of photosynthetically fixed C belowground (35–37). Some estimates suggest up to 40% of fixed C is translocated belowground (38), and grasses are suggested to be near that upper limit with ~30% of fixed C allocated belowground (39). These rhizodeposits also include root exudates which are composed of sugars, organic acids, phenolic compounds, and amino acids (1, 17, 40, 41). Differences in plant physiology influencing the quantity and composition of root exudates can affect rhizosphere bacterial community composition. For example, C4 grasses have higher photosynthetic rates (i.e., fix more C) and greater root biomass allocation compared to C3 plants, resulting in a greater quantity of root exudates (42, 43). C3 plant root exudates can contain a greater variety of organic acids and amino acids along with the sugars mannose, maltose, and ribose compared to C4 plant root exudates, which can contain several sugar alcohols (i.e., inositol, erythritol, and ribitol) (44). However, N fertilization has been shown to increase C assimilation in plants but decrease belowground allocation of assimilated C while increasing total C into soils as rhizodeposits. (39, 45). Prior studies revealed that root exudation of organic C can be higher in both low nutrient scenarios (46, 47) and high nutrient scenarios (48, 49). Further, differences in soil nutrient status can change the composition (i.e., carbohydrates, organic acids, and amino acid concentrations) of root exudates (46, 50). Thus, fertilization and plant specific rhizodeposition patterns of C3 forbs and C4 grasses are predicted to differentially affect rhizosphere bacterial community structure.

In this study, we address the following question: To what extent does long-term fertilization (N-P-K) of bulk soil shift rhizosphere bacterial communities of two plant species representing distinct functional types (i.e., a C3 forb and a C4 grass)? First, we hypothesize that nutrient addition will decrease bacterial species diversity and increase the abundance of copiotrophic taxa in all soils, especially rhizosphere soils due to increased availability of labile C from root exudates. We expect that fertilization will stimulate microbial activity of faster growing copiotrophic species, which would outcompete slower growing oligotrophic species and result in decreased bacterial diversity. This effect is predicted to be amplified within plant rhizospheres due to the availability of labile C substrates in root exudates, which should preferentially select for copiotrophic bacteria. Second, we hypothesize that fertilization will be the primary factor determining differences in rhizosphere communities and plant identity will secondarily influence the rhizosphere community. If bulk soil is the reservoir for the rhizosphere community, then fertilization will determine rhizosphere bacterial diversity and community composition more strongly. In addition, plant type can also affect rhizosphere communities due to differences in root exudate composition; however, fertilization effects will constrain rhizosphere effects. As a result, plant species are expected to associate with unique core microbiomes that differ between fertilization treatments.

To test these hypotheses, bulk and rhizosphere soils were sampled from two plant species (grass, forb) from fertilized and unfertilized plots at a long-term disturbance and fertilization experiment (established in 2003). Bacterial communities were identified using 16S rRNA amplicon sequencing which allowed binning of bacterial taxa as copiotrophic or oligotrophic by estimating the average rRNA *(rrn)* gene copy number. By evaluating differences in taxonomic information and 16S rRNA gene copy numbers of bulk and rhizosphere soils of two plant species with associated soil properties (i.e., ammonium, nitrate, soil pH, carbon, and moisture), we provide insight to biotic and abiotic processes that are contributing to rhizosphere bacterial community assembly.

## RESULTS

### Soil source and fertilization distinguishes soil properties

The main effect of fertilization was significantly different in the soil physiochemical property of pH (p=0.02); and the main effect of soil source (bulk vs. rhizosphere) was significantly different in the soil physiochemical properties of pH (p < 0.001), nitrate (p<0.0001), C percent (p=0.03), and N percent (p=0.04; Table S1). Rhizosphere soils were more similar to each other in soil properties than to bulk soils (Table 1, Tukey HSD, p<0.05). Specifically, bulk soil had lower total C and N, and nitrate concentrations than forb rhizospheres with grass rhizospheres having the highest values (Table 1, Tukey HSD, p<0.05). Soil pH was lowest in rhizosphere soils compared to bulk soils but higher in fertilized soils compared to unfertilized soils within soil sources (Table 1, Tukey HSD, p<0.05).

**Table 1.**
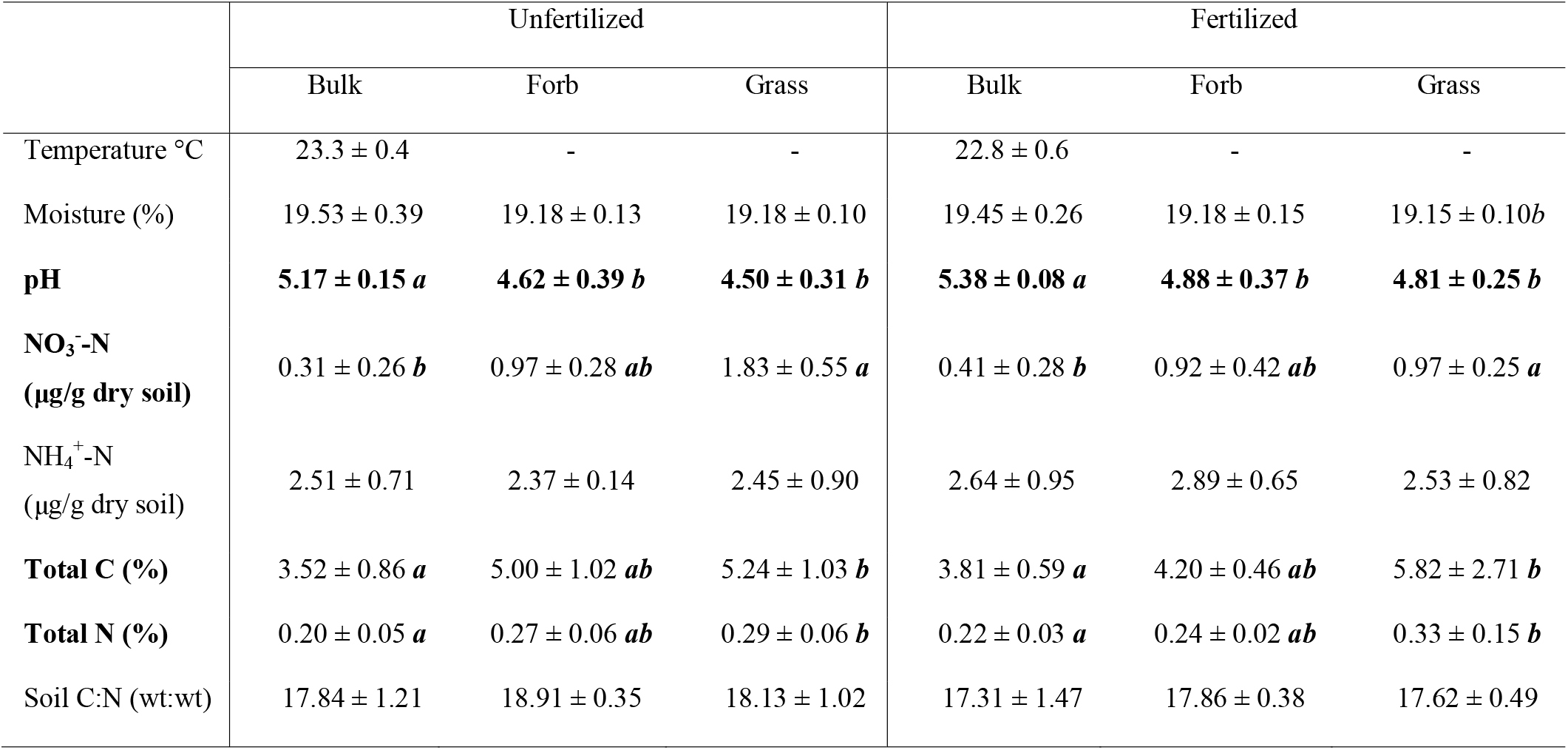
(A) Soil physiochemical properties after 12 years of fertilization and mowing disturbance. Average (mean ± SD) soil properties (temperature, gravimetric moisture, pH, extractable nitrate and ammonium concentrations, total soil C and N, and C:N ratio) across unfertilized and fertilized plots and among soil sources (bulk, forb rhizosphere, and grass rhizosphere). Fertilization main effect that is significantly differently (ANOVA p<0.05) is bolded. Letters represent significant differences between soil sources (Tukey’s HSD p < 0.05).

### Fertilization increased soil bacterial diversity in bulk and rhizosphere soils

Chao1 bacterial richness (p<0.0001) and Shannon H’ diversity (p<0.0001) were higher in fertilized soils compared to unfertilized soils (Table S2, Fig. 1A). In addition, the main effect of soil source influenced bacterial diversity; bulk soil bacterial diversity was significantly higher than rhizosphere soil diversity (Tukey HSD, p<0.05, Table S2, Fig. 1B). Finally, results revealed a positive relationship between Shannon H’ diversity and pH, where pH explained 71% (p=0.0003) and 32% (p=0.03) of the variation in bacterial diversity in unfertilized and fertilized treatments, respectively, across all soil sources (Fig. 1C).

**Figure 1.**
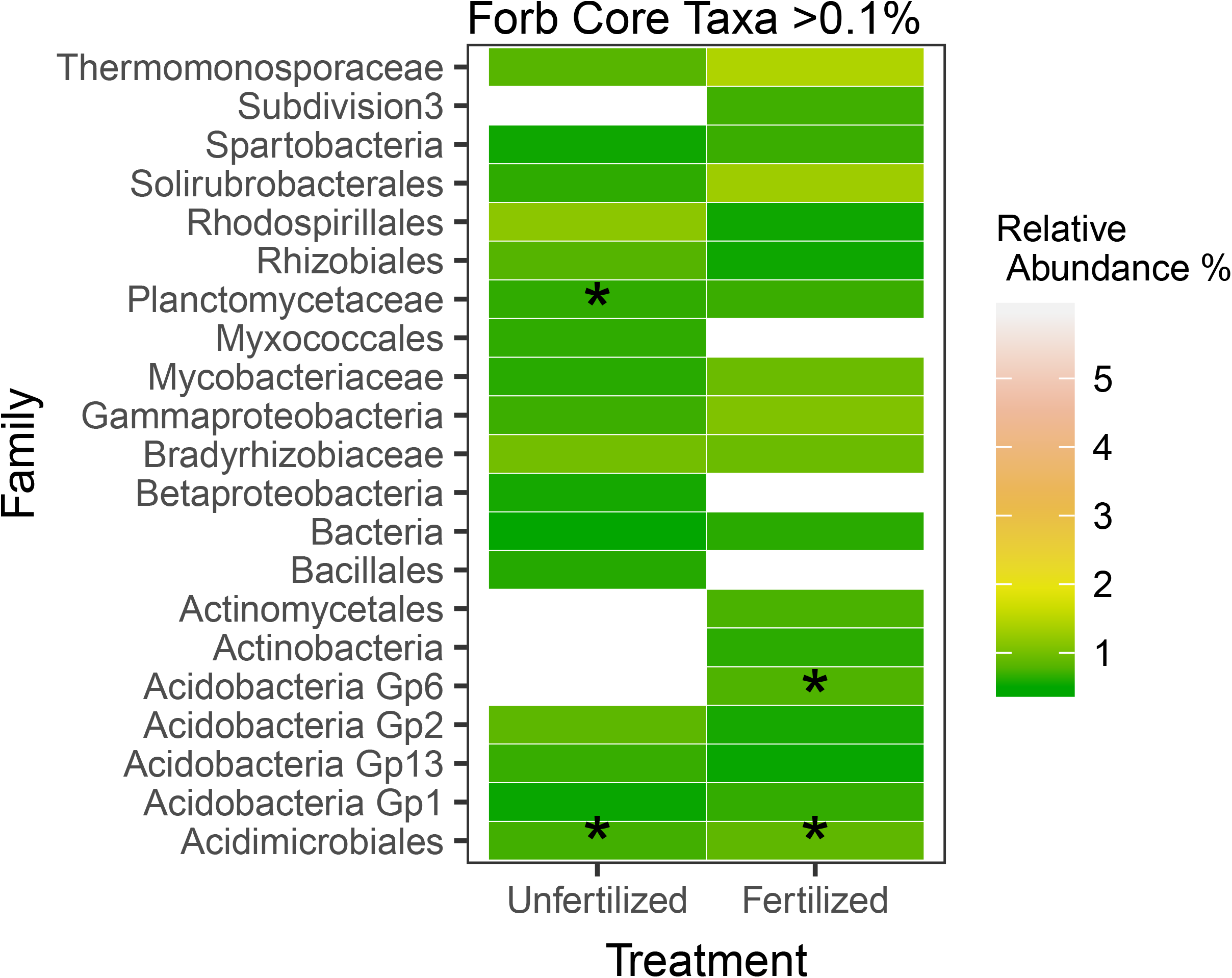
Bacterial diversity patterns according to soil source, fertilization, and soil pH. Boxplots of bacterial diversity for Chao1 richness (A) and Shannon H□ diversity index (B) associated with soil source (bulk, grass rhizosphere, forb rhizosphere) and fertilization treatment. Linear regression of soil pH and bacterial community Shannon H’ diversity by fertilization treatment with 95% confidence intervals (C); Fertilized: R^2^=0.32, p=0.03, Unfertilized: R^2^=0.71, p=0.0003.Colors indicate fertilization treatment (gray = unfertilized, green = fertilized) at mowed plots. Asterisks (*) indicate significant differences between fertilization treatments and letters represent significant differences between soil sources (Tukey’s HSD, p < 0.05).

### Copiotroph to oligotroph ratios indicated oligotroph-dominated bacterial communities

Across all samples we detected 9 to 30 copiotrophic and 82 to 190 oligotrophic taxa at the class level. This resulted in copiotroph to oligotroph ratios of < 0.2 within all treatment combinations. Nutrient additions significantly decreased the ratio of copiotrophs to oligotrophs in bulk soils compared to rhizosphere soils (Tukey’s HSD, p < 0.05; Table S4; Fig. 2). Finally, there was no relationship between bacterial Shannon H’ diversity and copiotroph to oligotroph ratio (Fertilized: R^2^=-0.01, p=0.38; Unfertilized: R^2^=0.14, p=0.13) (Fig. S1).

**Figure 2.**
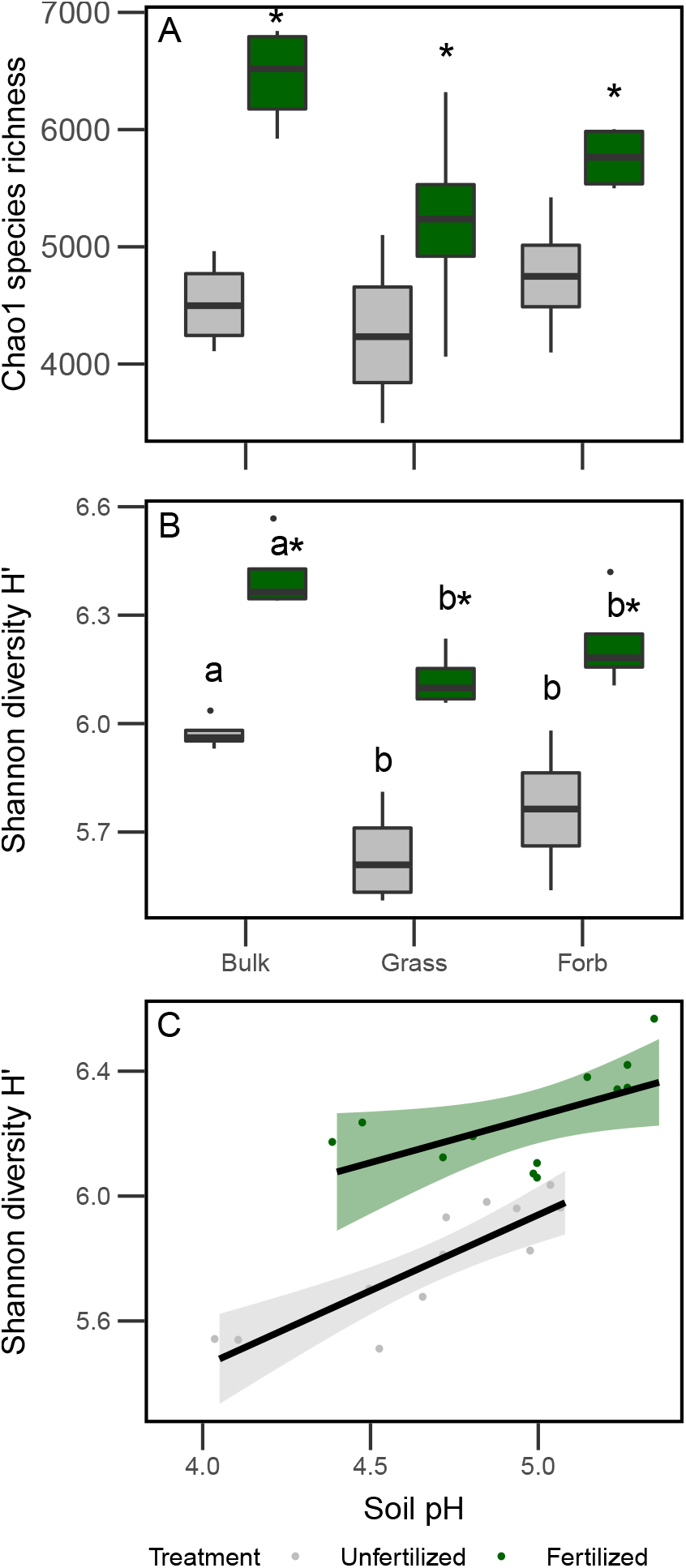
Comparison of bacterial life history traits. Boxplots of copiotroph to oligotroph ratios (based on 16S rRNA sequences) according to soil sources (bulk, grass rhizosphere, forb rhizosphere) and fertilization treatment. Boxplots are colored according to fertilization treatment (gray = unfertilized, green = fertilized). Letters indicate significant differences among soil sources (Tukey’s HSD, p < 0.05).

### Fertilization treatment and soil source influenced bacterial community composition

Specifically, fertilization treatment (along PCoA axis 1) explained 31.6% of variation in bacterial community composition, while soil source (primarily bulk vs. rhizosphere) separated bacterial composition (along PCoA axis 2) and explained 22.5% of bacterial community variation (Fig. 3). Main effects of soil source (PERMANOVA R^2^=0.23, P=0.001) and fertilization treatment (PERMANOVA R^2^=0.281, P=0.001) influenced bacterial community composition (Table S3A). According to pairwise comparisons, rhizosphere bacterial community composition was similar between grass and forb rhizosphere samples within fertilization treatments (Table S3B). When examining relationships between community composition and soil characteristics, higher soil pH and moisture were correlated to fertilized bulk soils (Fig. 3). Further, higher concentrations of soil C and N were correlated with rhizosphere community composition (Fig. 3).

**Figure 3.**
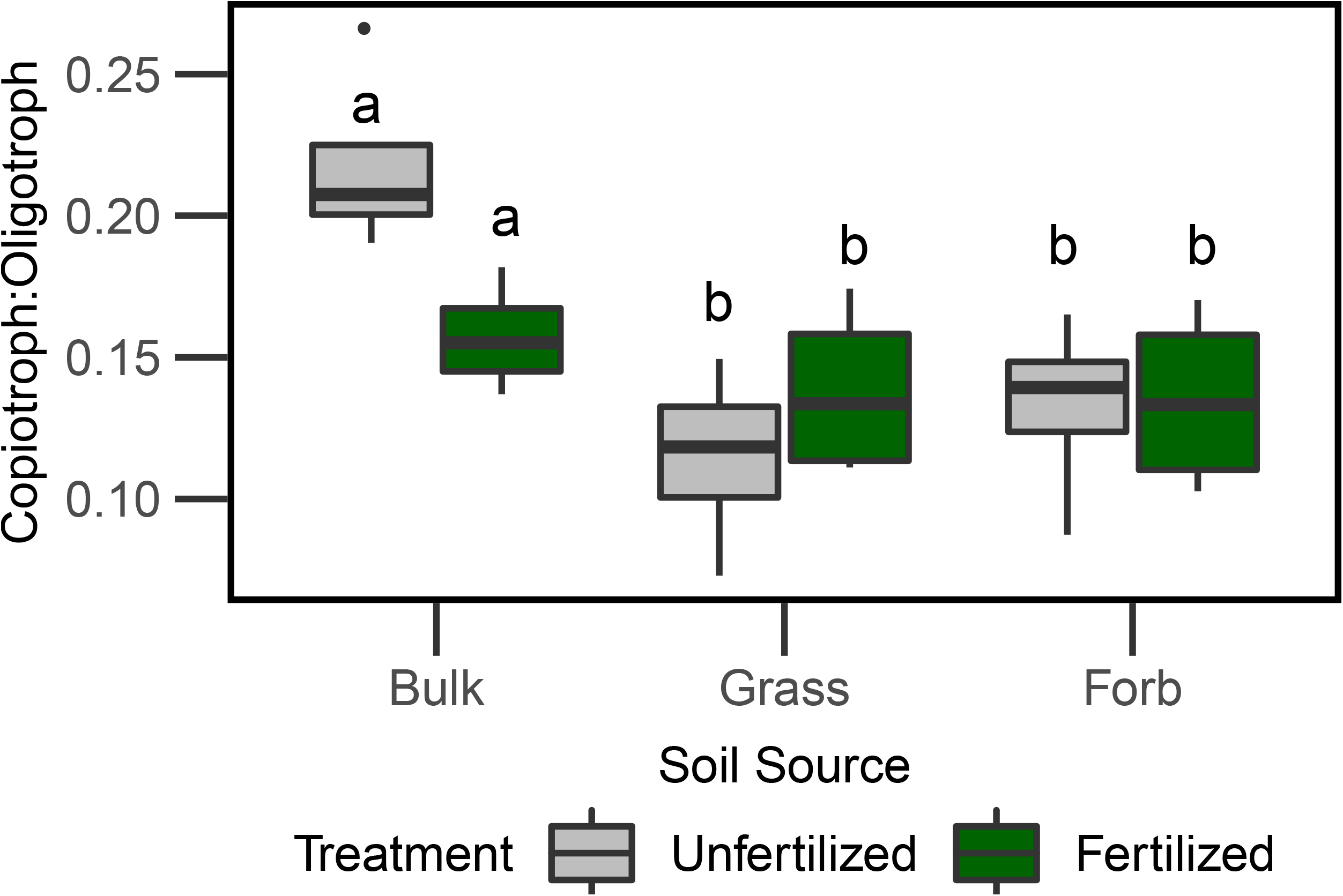
Ordination based on Principal Coordinates Analysis depicting bacterial community composition. Colors represent fertilization treatment (gray = unfertilized, green = fertilized) and symbols represent soil source (bulk soil = circle, grass rhizosphere = open square, forb rhizosphere = filled square). Vectors represent soil factors that are correlated to patterns in bacterial community composition (p<0.05) (pH = soil pH, moisture = soil gravimetric moisture percent, C% = total soil carbon, N%= total soil nitrogen).

### Different bacterial taxa (OTUs) represented fertilization treatments and plant species

We compared bacterial community taxonomic shifts in unfertilized and fertilized bulk soils and then grass and forb rhizospheres, concluding with differences in microbiome structure between the two plant species. Within bulk soil samples, important indicator species for bacterial communities within unfertilized plots were from the class Alphaproteobacteria with 1 OTU from the order Rhizobiales and 2 OTUs from Rhodospirillales and 3 OTUs from the class Spartobacteria (Table S5). In contrast, fertilized bulk soils were best represented by members of the class Actinobacteria with 1 OTU from the order Actinomycetales and 2 OTUs from the order Solirubrobacterales. While OTUs within Rhizobiales were identified as indicator species for bacterial communities in unfertilized bulk soils, this order was in greatest relative abundance compared to other orders within both fertilization treatments (Fig. S2).

Comparisons of rhizosphere bacterial OTU presence/absence data revealed that forb (1,249 OTUs) and grass (1,019 OTUs) rhizospheres have distinct but overlapping microbiomes. Of the 1,621 total OTUs found in rhizosphere soils, 647 are broadly-distributed and are observed in all plant rhizospheres and bulk soils regardless of treatment. Therefore, less than half of the forb (48%) and grass (37%) rhizosphere members were unique to that plant functional type, and broadly-distributed OTUs dominate plant microbiomes especially in grasses.

Of OTUs that were only represented in the grass microbiome (n=372), only 22 bacterial families are represented at > 0.075% relative abundance. Within those top OTUs, unfertilized grass rhizospheres were enriched in 9 families while fertilized plots were enriched in 19 families (Fig. 4). Indicator species for unfertilized grass rhizospheres included 2 OTUs, one in the genus *Singulisphaera* and family Planctomycetaceae (IndVal = 0.38, *P* = 0.026) and an unclassified Spartobacteria OTU (IndVal = 0.44, *P*=0.008; Table S5). Indicator species for fertilized grass rhizospheres included two OTUs, one in the genus *Planctomyces* and family Planctomycetaceae (IndVal = 0.42, *P*=0.011) and one in the genus *Actinoallomurus* and family Thermomonosporaceae (IndVal = 0.36, *P*=0.045; Table S5).

**Figure 4:**
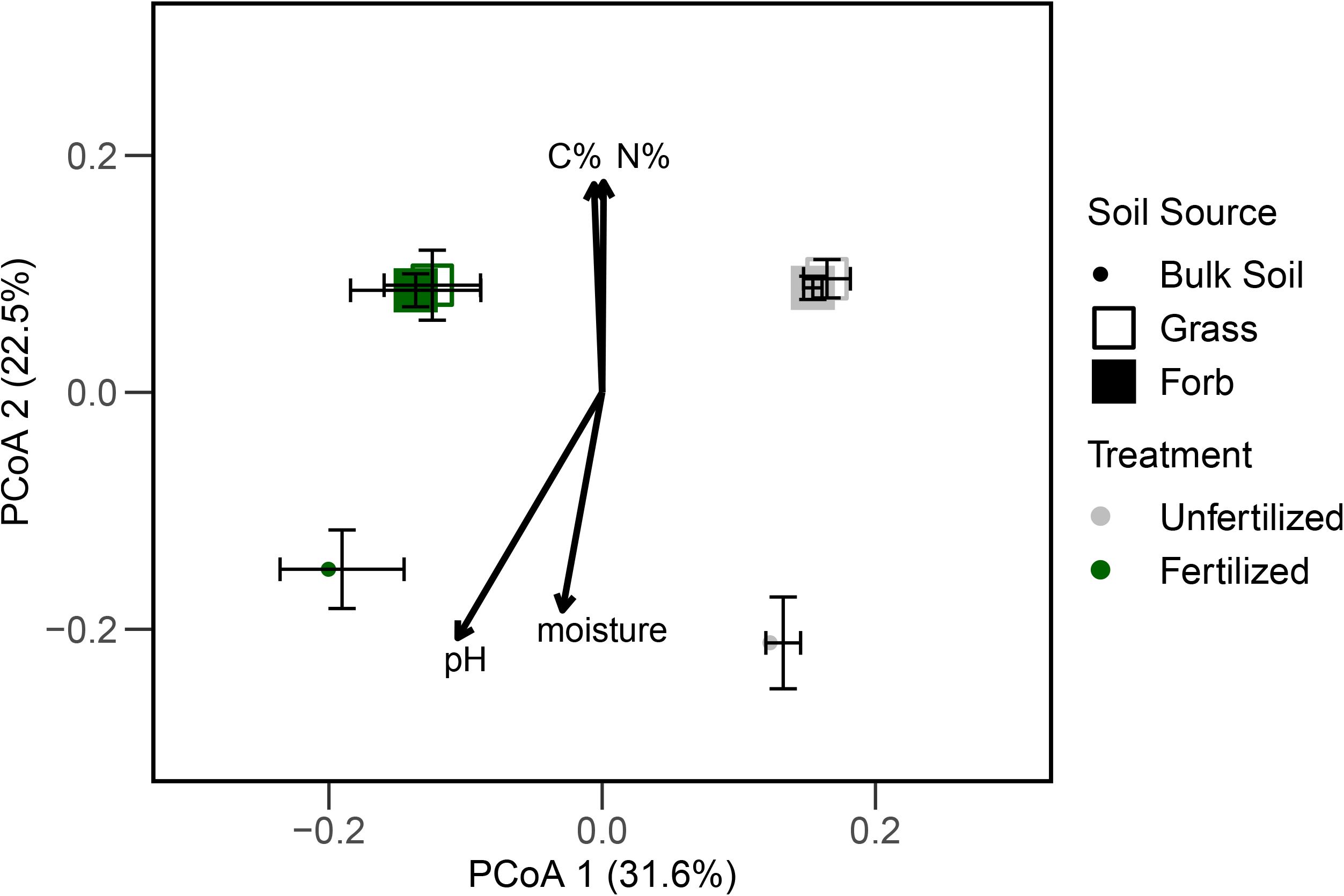
Comparisons of top OTU relative abundances (>0.075%) at the family level between fertilization treatments for grass rhizosphere bacterial communities. Asterisk (*) represents indicator species present within family (Table S5). Colors indicate relative abundance increases from cool to warm (green yellow, orange, and red). White boxes indicate taxa present at <0.075% relative abundance.

Of the OTUs that were only represented in the forb microbiome (n=602), only 21 bacterial families are represented at > 0.1% relative abundance. Within those top OTUs, unfertilized forb rhizospheres were enriched in 10 families while fertilized plots were enriched in 16 families (Fig. 5). Indicator species included two OTUs, Acidobacteria Gp1 (IndVal=0.42, *P*=0.02), and an unclassified Proteobacteria (IndVal=0.46, *P*=0.033; Table S5). Indicator species included an OTU in Acidobacteria Gp1 (IndVal= 0.34, *P*=0.041) class and an unclassified bacterial OTU (IndVal=0.60, *P*=0.017; Table S5).

**Figure 5:**
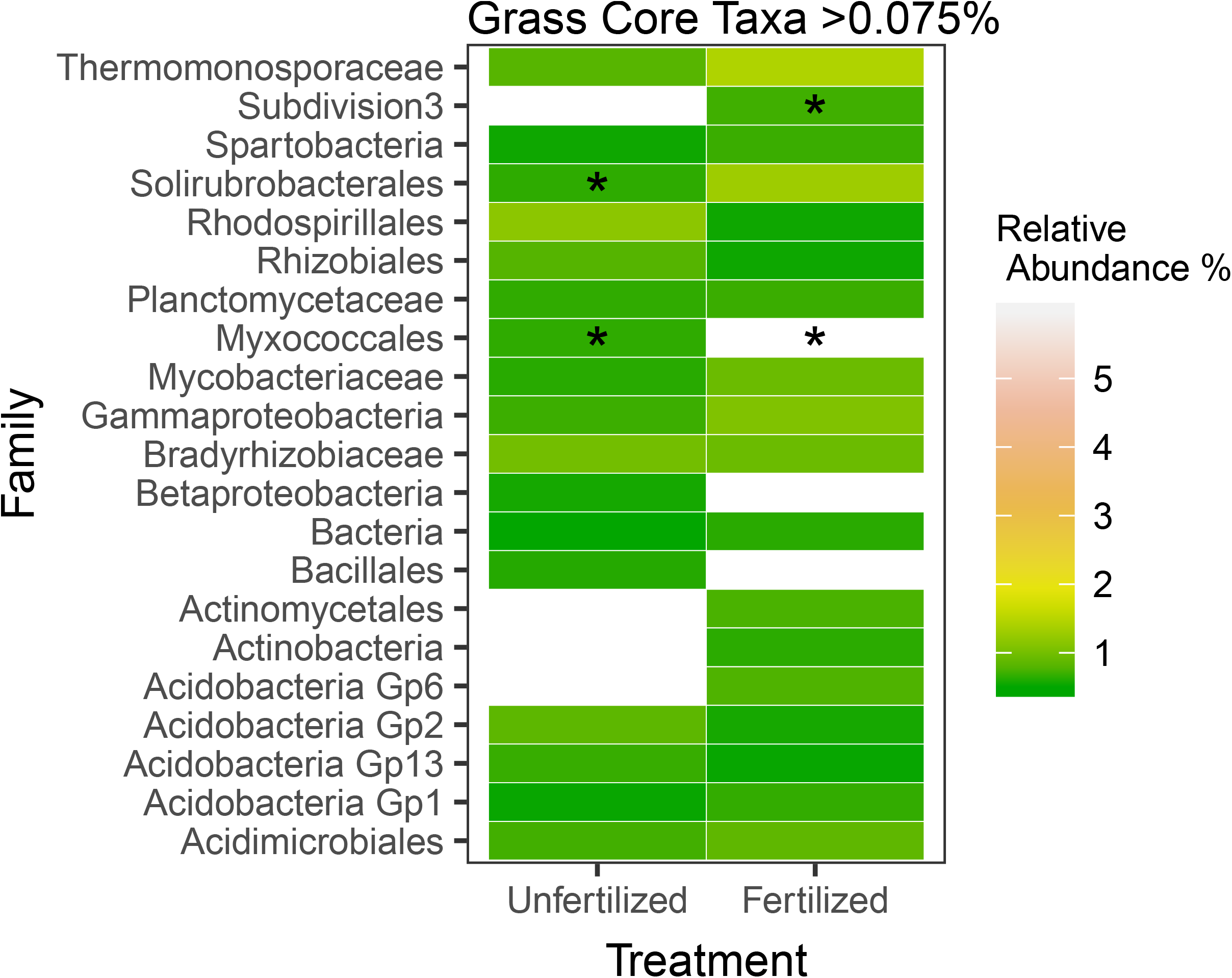
Comparisons of top OTU relative abundances (>0.1%) at the family level between fertilization treatments for forb rhizosphere bacterial communities. Asterisk (*) represents indicator species present within family (Table S5). Colors indicate relative abundance increases from cool to warm (green yellow, orange, and red). White boxes indicate taxa present at <0.1% relative abundance.

## Discussion

In this study, nutrient addition increased bacterial species diversity (H’) and richness in bulk and rhizosphere soils. These results were similar to O’Brien et al. (51) but contrary to our prediction and the results of other studies (14–16). Overall, bulk soils had the greatest bacterial diversity and highest pH values when compared to rhizosphere soils. Since pH is known to be a strong driver of bacterial diversity which can have a positive relationship to pH (52, 53), this increase in diversity may, in part, be due to the greater bulk soil pH compared to rhizosphere soil. The difference in pH between soil types is possibly due to organic acids in plant root exudates released into the rhizosphere (41), however, we did not analyze the composition of root exudates. Additionally, pH tended to be lower in unfertilized treatments, and diversity was more strongly related to pH in unfertilized soils compared to fertilized soils. This may be due to sensitivity of bacteria to acidic soils (53). The increase in bacterial diversity is likely the result of soil pH and niche differentiation due to fertilization increasing nutrient availability and rhizodeposition by plants, which introduces organic C resources for heterotrophs (17, 32). In dilution to extinction experiments, decreases in microbial diversity can result in loss of microbial functional diversity (54, 55). Therefore, increases in microbial diversity could result in increased microbial functional diversity, which could increase C cycling and promote N mining particularly in plant rhizospheres (56).

Bacterial taxa identified in rhizosphere samples are putatively involved in nutrient cycling and disease suppressive functions. For example, fertilized forb rhizospheres were enriched in taxa from the family Streptomycetaceae, of which many produce antibiotics (57) and Sphingomonadaceae, which include taxa with disease suppression potential against fungal pathogens (58) (Fig. 5). This increase in disease suppressive bacterial taxa suggest a potential increase in plant pathogenic taxa within fertilized rhizospheres; however, this study did not specifically address disease suppression in soils. In contrast, fertilized grass rhizospheres were enriched with taxa putatively involved in N2-fixation (Acetobacteraceae) (59) and also Chitiniphagaceae and Conexibacteraceae, which have been implicated in decomposition of recalcitrant C sources (60, 61) (Fig. 4). Bacterial taxa in the Xanthomonadaceae family, which have previously been found in environments containing glyphosate (62), and Caulobacteraceae, which grows optimally on pesticides (63), are also more abundant in fertilized grass rhizospheres (Fig. 4). Since fertilization increased bacterial diversity and shifted composition, it is possible that fertilization has stimulated root exudation. The relative increase in complex C degrading bacterial taxa in the grass rhizosphere could also be due greater inputs of phenolics and terpenoids used as allelochemicals by the plant as revealed in past studies (64, 65). These differences in bacterial composition between the two plants species could be due to differences in composition of root exudates released into the rhizosphere (37), however, we did not analyze the composition of root exudates in the present study. Together, results suggest that nutrient addition enriches forb rhizospheres with putatively disease suppressive bacteria and grass rhizospheres with taxa capable of decomposing complex C sources.

Within bulk soil bacterial members, putative nitrogen cycling taxa in the order Rhizobiales were enriched across all fertilization treatments (66, 67). This is not surprising considering the limited amount of nitrogen in both unfertilized and fertilized soils at the study site. Despite the increase in taxa capable of N2-fixation in fertilized rhizospheres, these bacteria will acquire soil N if it is available (68). Therefore, these taxa may be less cooperative with plant associates than the same taxa from unfertilized soils thereby reducing plant benefit (2, 69). This was not specifically tested in this study but could be an important future research topic.

Contrary to our prediction, bulk soils had a higher copiotroph to oligotroph ratio (based on *rrn* gene copy number) than rhizospheres. Characteristic of the copiotrophic life history strategy is the ability to rapidly decompose labile C sources, therefore we expected that C rich root exudates in the rhizosphere would support higher proportions of copiotrophic species (17). Additionally, fertilization did not increase the relative abundance of copiotrophic taxa. Rather, the observed copiotroph to oligotroph ratios were low in all samples with unfertilized bulk soils having the greatest proportion (22%) and unfertilized grass rhizospheres having the lowest (13%) copiotroph to oligotroph ratios. We suggest that the dominance of oligotrophs reflects the low-nutrient history of this wetland (29, 70), which is in contrast to agricultural systems that undergo regular fertilization at target rates intended to support high nutrient requirements for enhanced crop production (e.g., corn).

These results are in contrast to our first hypothesis and in agreement with our second hypothesis. Analyses of bacterial diversity and copiotroph to oligotroph ratios revealed an increase in bacterial diversity in response to fertilization and dominance of oligotrophs across all treatments within the study wetland. The low nutrient history of the study site is likely the primary factor shaping bacterial community composition within the wetland. In agreement with our second hypothesis, comparisons of bulk and rhizosphere bacterial communities revealed that rhizospheres were more similar to each other than to bulk soil bacterial communities within fertilization treatments. Core plant microbiomes were predominantly composed of broadly-distributed taxa; therefore, changes in bulk soil bacterial composition due to nutrient enrichment can directly alter plant microbiome composition and indirectly diminish benefits to plants if nutrient enrichment selects for more competitive bacterial taxa. These results highlight the importance of bulk soils as reservoirs of diversity for plant rhizospheres, which could have further implications for agricultural plant species in maintaining beneficial microbial communities.

Overall, this study revealed that long-term fertilization of oligotroph-dominated soils in low nutrient wetlands increases bacterial species diversity. This increase in bacterial diversity has the potential to result in increased C and nutrient cycling that could lead to declines of wetland C storage potential. Nutrient enrichment also differentially alters plant rhizosphere composition in a way that suggests metabolic changes within soil bacterial communities. These metabolic changes could indirectly impact plant species diversity by providing an advantage to one species versus another through disease suppression or by increasing plant available N through promotion of soil organic matter decomposition. If indirect fertilization supports rhizosphere bacterial communities that can enhance recalcitrant or labile C decomposition, wetland C storage potential could decline. Based on this study, bacterial taxonomic characterization sheds light on fertilization effects on plant-bacterial relationships. As such, nutrient enrichment effects on the metabolic diversity of bacterial communities could be even more pronounced and warrants further investigation.

## Material and Methods

### Study site and experimental design

A long-term experimental site established in 2003 to test the effects of fertilization, mowing, and the interaction on wetland plant communities. The site is located at East Carolina University’s West Research Campus in Greenville, North Carolina, USA (35.6298N, −77.4836W). A description of the study site and experimental design can be found in Goodwillie and Franch (71) and is summarized here. This site is classified as a jurisdictional wetland but historically described as a mosaic of wet pine flatwood habitat, pine savanna, and hardwood communities. Soils were characterized as fine, kaolinitic, thermic Typic Paleaquults (Coxville series) with a fine sandy loam texture which are ultisols that are acidic and moderate to poorly drained soil types (https://soilseries.sc.egov.usda.gov/osdname.aspx). The annual mean temperature is 17.2 °C and annual precipitation is 176 cm (https://www.climate.gov/maps-data/dataset/). Treatments are replicated on eight 20×30 m blocks, and the N-P-K 10-10-10 pellet fertilizer is applied 3× per year (February, June, and October) for a total annual supplementation of 45.4 kg ha^-1^ for each nutrient. Plots are mowed by bush-hog and raked annually to simulate a fire disturbance (71).

We compared rhizosphere and bulk soil microbiomes in mowed unfertilized and fertilized plots, where herbaceous species dominated. Soil samples were collected at mowed/unfertilized and mowed/fertilized plots in four out of eight replicate blocks to reduce variability due to hydrology. Half the site is located adjacent to a ditch (drier soils) compared to away from the ditch, where soil conditions are wetter. Since this hydrologic gradient has resulted in distinct plant communities (C. Goodwillie M.W. McCoy and A. L. Peralta, submitted for publication), we collected samples from the wetter plots (away from the drainage ditch).

### Bulk and rhizosphere soil sampling

We collected soil samples on September 29, 2015, approximately three months after last fertilization treatment. Due to annual mowing and raking in sample plots, there was limited biomass accumulated in the organic horizon. We focused soil sampling and analysis on the mineral horizon. For a single composite bulk soil sample, we collected two soil cores (12 cm depth, 3.1 cm diameter) near each of the three permanently installed 1 m^2^ quadrats used for annual plant surveys. Each composite bulk soil sample was homogenized, passed through a 4 mm sieve, and any plant material removed before further analysis. At each plot, rhizosphere soils were collected from the C3 forb *Euthamia caroliniana* (L.) Greene ex Porter & Britton and C4 grass *Andropogon virginicus L.* Rhizosphere soils were a composite of three root systems of the same species. Roots were gently dislodged from soil and neighboring roots and placed in a paper bag. After vigorous shaking, soil in the bag was processed for abiotic analysis. The roots were placed into 50 mL centrifuge tubes with 30 mL sterilized Nanopure^®^ water and shaken at 100 RPM for 1 hour. Washed roots were removed, and the soil and water mixture was freeze-dried to remove water. Freeze-dried rhizosphere samples were stored at −80 °C until DNA extraction.

### Soil chemical and physical characteristics

We measured gravimetric soil moisture by drying 20-30 g of field-moist soil at 105 °C for 24 hours. We calculated percent moisture as the difference in weight of moist and dried soils divided by the oven-dried soil weight. Oven-dried samples were ground and measured for pH by mixing a 1:1 (soil:water) solution. A subsample of oven-dried soil was sieved with a 500 μm mesh and analyzed for total carbon and total nitrogen (TC, TN) using an elemental analyzer (2400 CHNS Analyzer; Perkin Elmer; Waltham, Massachusetts, USA) at the Environmental and Agricultural Testing Service laboratory (Department of Crop and Soil Sciences at NC State). Approximately 5 g of field moist soil was extracted with 45 ml of 2 M KCl, and available ammonium (NH_4_^+^) and nitrate (NO_3_^-^) ions were colorimetrically measured using a SmartChem 200 auto analyzer (Unity Scientific Milford, Massachusetts, USA) at the East Carolina University Environmental Resources Laboratory.

### Bacterial community analyses

We extracted DNA from soils using the Qiagen DNeasy PowerSoil Kit. We used this DNA as template in PCR reactions using barcoded primers (bacterial/archaeal 515FB/806R) originally developed by the Earth Microbiome Project to target the V4 region of the bacterial 16S subunit of the ribosomal RNA gene (72). For each sample, three 50 μL PCR libraries were prepared by combining 30.75 μL molecular grade water, 5 μL Perfect Taq 10x buffer, 10 μL Perfect Taq 5x buffer, 1 μL dNTPs (40 mM total, 10 mM each), 0.25 μL Perfect Taq polymerase, 1 μL forward barcoded primer (10 μM), 1 μL reverse primer (10 μM), and 1 μL DNA template (10 ng μL^-1^). Thermocycler conditions for PCR reactions were as follows: initial denaturation (94 °C for 3 minutes); 30 cycles of 94 °C for 45 seconds, 50 °C for 30 seconds, 72 °C for 90 seconds; final elongation (72 °C, 10 minutes). Triplicate PCR reactions were combined and cleaned using the AMPure XP magnetic bead protocol (Axygen, Union City, California, USA). Cleaned PCR product were quantified using QuantIT dsDNA BR assay (Thermo Scientific, Waltham, Massachusetts, USA) and diluted to a concentration of 10 ng μL^-1^ before pooling libraries in equimolar concentration of 5 ng μL^-1^. We sequenced pooled libraries using the Illumina MiSeq platform using paired end reads (Illumina Reagent Kit v2, 500 reaction kit) at the Indiana University Center for Genomics and Bioinformatics Sequencing Facility. Sequences were processed using mothur (v1.40.1) (73) MiSeq pipeline (74). We assembled contigs from the paired end reads, quality trimmed using a moving average quality score (minimum quality score 35), aligned sequences to the SILVA rRNA database (v128) (75), and removed chimeric sequences using the VSEARCH algorithm (76). We created operational taxonomic units (OTUs) by first splitting sequences based on taxonomic class and then binning into OTUs based on 97% sequence similarity. The SILVA rRNA database (v128) (75) was then used to assign taxonomic designations to OTUs.

Samples were rarefied to 43,811 OTUs and resampled. We used *vegan::diversity* (77) to calculate bacterial species diversity as calculating Shannon diversity index (H’) because it accounts for species abundance and evenness and rare species (78, 79). We estimated bacterial richness using Chao1 species richness because it is non-parametric and also considers rare species (79, 80). Shannon diversity was calculated using the *vegan::diversity* function and Chao1 OTU richness using *vegan::estimate* (77). We assigned gene copy number to each OTU using RDP classifier (v2.12) (81) integrated with the *rrn* operon database developed by the Schimdt

Laboratory at the Michigan Microbiome Project, University of Michigan (23, 27). Higher gene copy numbers (>=5) represent the copiotrophic lifestyle and lower gene copy numbers (<5) represent the oligotrophic lifestyle (20, 24, 82). The number of copiotrophs and oligotrophs were summed for each soil sample to calculate the copiotroph to oligotroph ratio within a soil bacterial community.

### Statistical Analyses

All statistical analyses were performed in the R statistical environment (RStudio v1.1.383, Rv3.4.0) (83). We used two-way model of analysis of variance (ANOVA) to compare main effects of soil source and fertilization treatment and the interaction to test for differences in OTU diversity and richness, copiotroph to oligotroph ratios, and soil parameters (soil pH, total carbon, extractable ammonium and nitrate total nitrogen, soil moisture). Significant interactions were compared with Tukey’s post-hoc analysis using the *agricolae::HSD.test* R function (84). We examined diversity by visualizing bacterial community responses to fertilization and rhizosphere association using principal coordinates of analysis (PCoA) based on Bray-Curtis dissimilarity. We used permutational multivariate analysis of variance (PERMANOVA) to test for differences in bacterial community composition among treatments and within treatment using pairwise comparisons. Hypothesis testing using PERMANOVA was performed using the *vegan::adonis* function (77). We examined the relationship between soil parameters and bacterial Bray-Curtis dissimilarity patterns using the *vegan::envfit* function (77). Soil parameters with p<0.05 were represented on the PCoA plot as vectors scaled by strength of correlation. We performed Dufrene-Legendre indicator species analysis using the *labdsv::indval* function (85) to identify specific community members that represented each soil source and fertilization treatment combination.

## Abbreviations

C: carbon
N: nitrogen
OTU: operational taxonomic unit
P: phosphorus
K: potassium

## Author contributions

RBB, CG, and ALP conceived and designed the research; RBB collected and analyzed the data; RBB wrote the manuscript; all authors performed field work and edited the manuscript.

## Acknowledgments

We thank M. Beamon, C. Eakins, J. LeCrone, J. Stiller, and S. Wilkinson for laboratory and field assistance. We thank J. Gill and the East Carolina University grounds crew for their efforts in maintaining the long-term ecological experiment. This work was supported by the National Science Foundation (GRFP to RBB and DEB 1845845 to ALP) and East Carolina University.

We also thank two anonymous reviewers for their suggestions that greatly improved this manuscript. All code and data used in this study can be found in a public GitHub repository (https://github.com/PeraltaLab/WRC15_Rhizo) and the NCBI SRA (BioProject PRJNA599142).

## Supplementary Material

**Supplemental Table S1.** Summary of two-way ANOVA comparing soil properties among soil source (bulk, grass rhizosphere, forb rhizosphere) and fertilization treatments.

**Supplemental Table S2.** Summary of two-way ANOVA comparing bacterial community Chao1 richness (A) and Shannon H’ diversity (B) metrics among soil source and fertilization treatments. Source represents bulk, grass rhizosphere, and forb rhizosphere and treatment represents fertilized and unfertilized mowed treatments. Main effects that were significantly different (ANOVA p<0.05) are bolded.

**Supplemental Table S3**. Summary of PERMANOVA main effects (soil source and fertilization treatment) and interaction (A) and pairwise PERMANOVA comparisons of soil sources (bulk, grass rhizosphere, forb rhizosphere) within fertilization treatments (B).

**Supplemental Table S4**. Summary of two-way ANOVA comparing bacterial community copiotroph to oligotroph ratio among soil source and fertilization treatments.

**Supplemental Figure S1:** Linear regression of copiotroph to oligotroph ratio and Shannon diversity H’ by fertilization treatment. Gray confidence bands represent 95% confidence intervals. Fertilized: R^2^=-0.01, p=0.38; Unfertilized: R^2^=0.14, p=0.13.

**Supplemental Figure S2:** Comparisons of bulk soil top OTU relative abundances (>1%) grouped by Order. Single asterisk (*) = indicator taxa for unfertilized treatment and double asterisk (**) = indicator taxa for fertilized plots (Table S5). Boxplots are colored according to fertilization treatment (gray = unfertilized, green = fertilized).

**Supplemental Table S5**: Summary of bacterial taxa (OTUs) characteristic to each soil source and fertilization treatment based on indicator species analysis. Listed are the top OTUs that are significantly associated with each soil source and fertilization treatment group.

