## Supplemental Data for "Long-term nutrient enrichment of an oligotroph-dominated wetland increases bacterial diversity in bulk soils and plant rhizospheres"

**Table S1.** Soil properties two-way ANOVA main effects table comparing soil sources (bulk, grass rhizosphere, forb rhizosphere) among treatments.

|  | Treatment | | Source | | T*S | |
| --- | --- | --- | --- | --- | --- | --- |
|  | F-Value | P | F-Value | P | F-Value | P |
| **Moisture (%)** | 0.28 | 0.84 | **5.14** | **0.02** | 0.05 | 0.95 |
| **pH** | 2.11 | 0.13 | **12.14** | **<0** | 0.07 | 0.93 |
| **NO3--N (μg/g dry soil)** | 0.81 | 0.51 | **12.51** | **<0** | 3.23 | 0.06 |
| NH4+-N (μg/g dry soil) | 0.25 | 0.86 | 0.07 | 0.93 | 0.21 | 0.81 |
| **Total C (%)** | 0.33 | 0.81 | **4.37** | **0.03** | 0.54 | 0.59 |
| **Total N (%)** | 0.27 | 0.85 | **3.96** | **0.04** | 0.42 | 0.66 |
| Soil C:N (wt:wt) | 1.37 | 0.28 | 1.54 | 0.24 | 0.22 | 0.81 |

**Table S2.** Summary of two-way ANOVA comparing bacterial community Chao1 richness (A) and Shannon H’ diversity (B) metrics. Source represents bulk, grass rhizosphere, and forb rhizosphere and treatment represents fertilized and unfertilized mowed treatments. Bolded values are considered significant.

(A) Chao1 richness

| Fixed Effect | SumSq | MeanSq | NumDF | F value | Pr(>F) |
| --- | --- | --- | --- | --- | --- |
| Source | 2323441 | 1161721 | 2 | 3.40 | 0.056 |
| **Treatment** | 10062645 | 10062645 | 1 | 29.476 | **<0.0001** |
| Source:treatment | 202762 | 101381 | 2 | 1.79 | 0.195 |

(B) Shannon diversity

| Fixed Effect | SumSq | MeanSq | NumDF | F value | Pr(>F) |
| --- | --- | --- | --- | --- | --- |
| **Source** | 0.399 | 0.199 | 2 | 12.901 | **0.0003** |
| **Treatment** | 1.278 | 1.278 | 1 | 82.705 | **<0.0001** |
| Source:treatment | 0.003 | 0.015 | 2 | 0.082 | 0.922 |

**Table S3**. (A) PERMANOVA main effects (soil source and fertilization treatment) and interaction, (B) pairwise PERMANOVA comparisons of soil sources (bulk, grass rhizosphere, forb rhizosphere) within fertilization treatments and (C) pairwise PERMANOVA comparisons of soil sources between fertilization treatments.

(A) Main effects

|  | SumSq | F-value | R^2^ | p-value |
| --- | --- | --- | --- | --- |
| **Source** | 0.466 | 4.924 | 0.234 | **0.001** |
| **Treatment** | 0.558 | 11.80 | 0.281 | **0.001** |
| Source * Treatment | 0.114 | 1.202 | 0.057 | 0.257 |

(B) Pairwise PERMANOVA within fertilization treatments

|  | Unfertilized | | | | Fertilized | | | |
| --- | --- | --- | --- | --- | --- | --- | --- | --- |
| Soil Sources | SumSq | F-value | R^2^ | p-value | SumSq | F-value | R^2^ | p-value |
| **Bulk x Forb** | 0.214 | 4.839 | 0.446 | **0.033** | 0.189 | 2.987 | 0.332 | **0.024** |
| **Bulk x Grass** | 0.215 | 5.123 | 0.461 | **0.034** | 0.169 | 3.011 | 0.334 | **0.036** |
| Forb x Grass | 0.035 | 0.819 | 0.120 | 0.557 | 0.072 | 1.223 | 0.169 | 0.186 |

**Table S4**. Main effects summary of two-way ANOVA comparing bacterial community copiotroph to oligotroph ratio.

| Fixed Effect | SumSq | MeanSq | NumDF | F value | Pr(>F) |
| --- | --- | --- | --- | --- | --- |
| **Source** | 0.757 | 0.379 | 2 | 7.257 | **0.005** |
| Treatment | 0.007 | 0.007 | 1 | 0.136 | 0.717 |
| Source:treatment | 0.939 | 0.142 | 2 | 2.716 | 0.093 |

**Figure S1**: Linear regression of copiotroph to oligotroph ratio and Shannon diversity H’ by fertilization treatment with 95% confidence intervals. R^2^=-0.011, p=0.40.

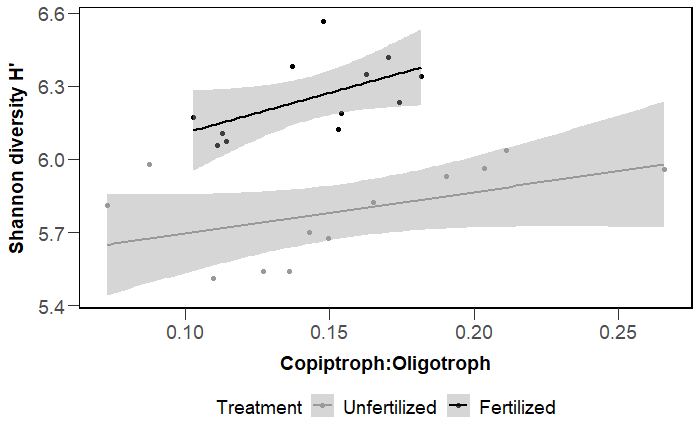

**Figure S2:** Comparisons of top OTU relative abundances (>1%) of bulk soils. Single asterisk (*) = indicator taxa for unfertilized treatment and double asterisk (**) = indicator taxa for fertilized plots (Table S#). Boxplots are colored according to fertilization treatment (gray = unfertilized, green = fertilized).

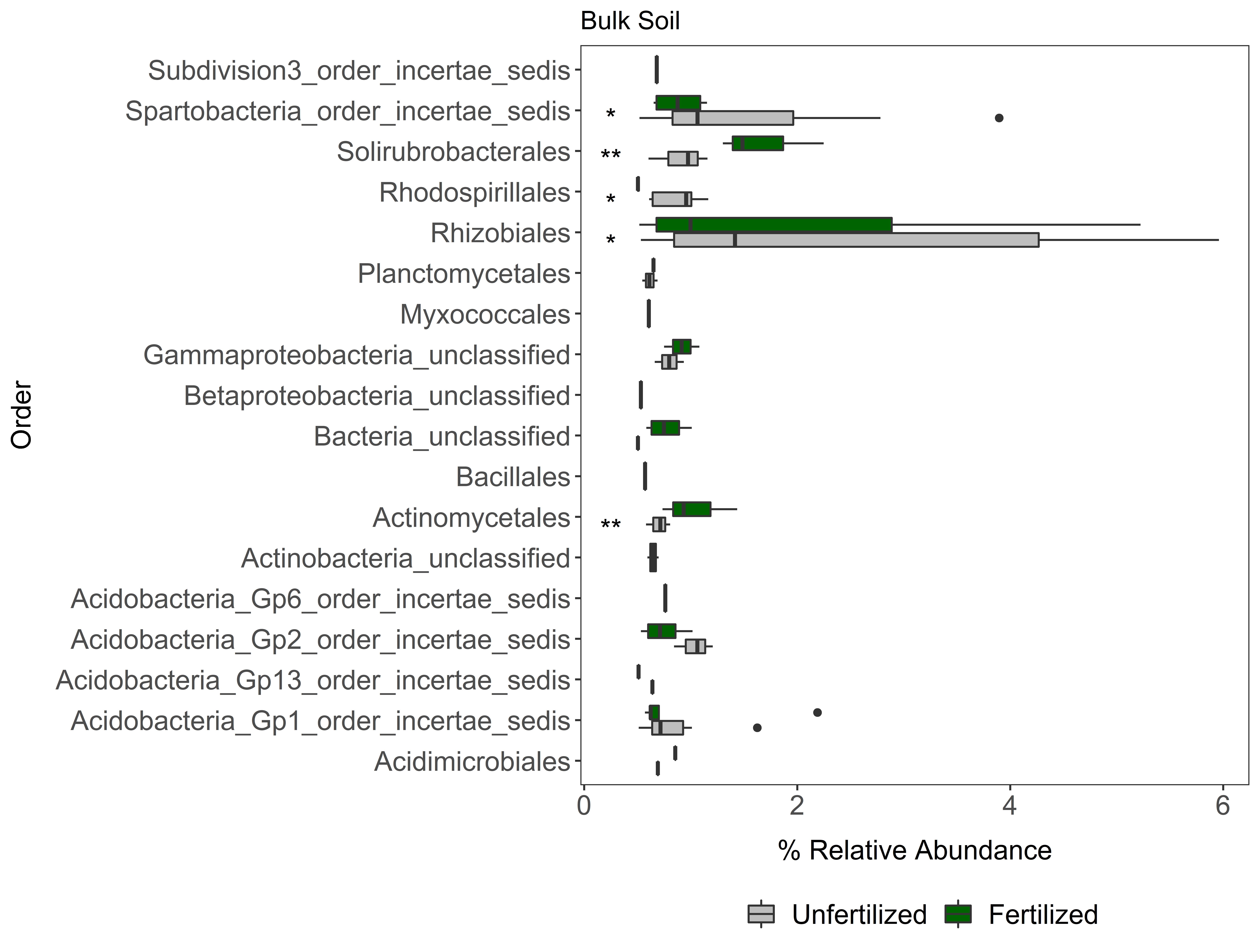

**Table S5**: Indicator species analysis output.

| OTU | Cluster | IndVal | Prob | Classification Phylum; Class; Order; Family; Genus |
| --- | --- | --- | --- | --- |
| Otu00016 | fertilized bulk | 0.716214922 | 0.028 | Actinobacteria; Actinobacteria; Solirubrobacterales; Solirubrobacterales_unclassified; Solirubrobacterales_unclassified |
| Otu00005 | fertilized bulk | 0.697698663 | 0.038 | Actinobacteria; Actinobacteria; Solirubrobacterales; Solirubrobacterales_unclassified; Solirubrobacterales_unclassified |
| Otu00013 | fertilized bulk | 0.640263156 | 0.03 | Actinobacteria; Actinobacteria; Actinomycetales; Thermomonosporaceae; Actinoallomurus |
| Otu00001 | unfertilized bulk | 0.592212461 | 0.022 | Proteobacteria; Alphaproteobacteria; Rhizobiales; Rhizobiales_unclassified; Rhizobiales_unclassified |
| Otu00039 | unfertilized bulk | 0.763467141 | 0.03 | Proteobacteria; Alphaproteobacteria; Rhodospirillales; Rhodospirillales_unclassified; Rhodospirillales_unclassified |
| Otu00029 | unfertilized bulk | 0.713504849 | 0.034 | Proteobacteria; Alphaproteobacteria; Rhodospirillales; Rhodospirillales_unclassified; Rhodospirillales_unclassified |
| Otu00064 | unfertilized bulk | 0.768348523 | 0.034 | Proteobacteria; Alphaproteobacteria; Rhodospirillales; Rhodospirillales_unclassified; Rhodospirillales_unclassified |
| Otu00018 | unfertilized bulk | 0.723635002 | 0.03 | Verrucomicrobia; Spartobacteria; Spartobacteria_order_incertae_sedis; Spartobacteria_family_incertae_sedis; Spartobacteria_genera_incertae_sedis |
| Otu00010 | unfertilized bulk | 0.64221502 | 0.033 | Verrucomicrobia; Spartobacteria; Spartobacteria_order_incertae_sedis; Spartobacteria_family_incertae_sedis; Spartobacteria_genera_incertae_sedis |
| Otu00003 | unfertilized bulk | 0.706878811 | 0.035 | Verrucomicrobia; Spartobacteria; Spartobacteria_order_incertae_sedis; Spartobacteria_family_incertae_sedis; Spartobacteria_genera_incertae_sedis |
| Otu00026 | fertilized forb | 0.341170952 | 0.041 | Acidobacteria; Acidobacteria_Gp1; Acidobacteria_Gp1_order_incertae_sedis; Acidobacteria_Gp1_family_incertae_sedis; Gp1 |
| Otu00023 | fertilized forb | 0.595088378 | 0.017 | Bacteria_unclassified; Bacteria_unclassified; Bacteria_unclassified; Bacteria_unclassified; Bacteria_unclassified |
| Otu00044 | unfertilized forb | 0.418012971 | 0.02 | Acidobacteria; Acidobacteria_Gp1; Acidobacteria_Gp1_order_incertae_sedis; Acidobacteria_Gp1_family_incertae_sedis; Gp1 |
| Otu00034 | unfertilized forb | 0.455001619 | 0.033 | Proteobacteria; Proteobacteria_unclassified; Proteobacteria_unclassified; Proteobacteria_unclassified; Proteobacteria_unclassified |
| Otu00027 | fertilized grass | 0.415830599 | 0.011 | Planctomycetes; Planctomycetacia; Planctomycetales; Planctomycetaceae; Planctomyces |
| Otu00013 | fertilized grass | 0.358524056 | 0.045 | Actinobacteria; Actinobacteria; Actinomycetales; Thermomonosporaceae; Actinoallomurus |
| Otu00024 | unfertilized grass | 0.38410856 | 0.026 | Planctomycetes; Planctomycetacia; Planctomycetales; Planctomycetaceae; Singulisphaera |
| Otu00003 | unfertilized grass | 0.441870014 | 0.008 | Verrucomicrobia; Spartobacteria; Spartobacteria_order_incertae_sedis; Spartobacteria_family_incertae_sedis; Spartobacteria_genera_incertae_sedis |
